# Sex differences in the behavioral and synaptic consequences of a single exposure to cannabinoid at puberty and adulthood

**DOI:** 10.1101/415067

**Authors:** Milene Borsoi, Antonia Manduca, Anissa Bara, Olivier Lassalle, Anne-Laure Pelissier-Alicot, Olivier J. Manzoni

## Abstract

Heavy cannabis consumption among adolescents is associated with significant and lasting neurobiological, psychological and health consequences that depend on the age of first use. Chronic exposure to cannabinoid (CB) agonists during adolescence alters social behavior and prefrontal cortex (PFC) activity in adult rats. However, sex differences on social behavior as well as PFC synaptic plasticity after acute CB activation remain poorly explored. Here, we determined the consequences of a single CB activation differently affects PFC in males and females by assessing social behavior and PFC neuronal and synaptic functions in rats during pubertal or adulthood periods, 24h after a single in-vivo cannabinoid exposure (SCE). During puberty, SCE reduced play behavior in females but not males. In contrast, SCE impaired sociability in both sexes at adulthood. General exploration and memory recognition remained normal at both ages and both sexes. At the synaptic level, SCE ablated endocannabinoid-mediated long-term depression (eCB-LTD) in the PFC of females of both ages and heightened excitability of PFC pyramidal neurons at adulthood, while males were spared. In contrast, SCE was associated to impaired long-term potentiation in adult males. Together, the data indicate behavioral and synaptic sex differences in response to a single in-vivo exposure to cannabinoid at puberty and adulthood.

## Introduction

Cannabis is the most frequently and widely used illicit drug among adolescents in developed countries (Gowing et al., 2015). Heavy cannabis consumption among adolescents is associated with significant and lasting neurobiological, psychological and health consequences developing in a dose-dependent fashion which are influenced by age of first use (Lisdahl et al., 2018; Iede et al., 2017; Levine et al., 2017). Chronic adolescent exposure to cannabinoids is linked to persistent adverse effects such as poor cognitive and psychiatric outcomes in adulthood (Levine et al., 2017) and regular cannabis use is associated with psychosocial impairment even in users without cannabis use disorder (Foster et al., 2017).

The primary psychoactive compound of the plant *Cannabis sativa*, Δ-9-tetrahydrocannabinol (THC), as well as the main endogenous cannabinoids (eCB) anandamide and 2-arachidonoylglycerol, all engage the same primary target in the central nervous system: the G-protein coupled cannabinoid receptor type 1 (CB1R). The eCB system consists of this and other receptors, eCB, and the enzymatic machinery for eCB synthesis and degradation (Hu and Mackie, 2015) and participates in neuronal development and synaptic plasticity in most brain areas (Gaffuri et al., 2012; Manduca et al., 2012; Lu and Mackie, 2016).

Adolescence is a period of profound morphological, neurodevelopmental and behavioral maturation. Brain volumes, sex steroids, and cortical morphometry all contribute to sex influences on developmental trajectories which are accompained by changes in the behavioral repertoire normally observed in this transitional period from infancy to adulthood. Puberty is characterized by external physical signs and hormonal alterations whose onset is signaled by gonadotropin-releasing hormone (Harris and Levine, 2003; Ojeda et al., 2003; Spear, 2000). This period is elicited through the complex interaction of endogenous and environmental factors (Sisk and Foster, 2004). Both adolescence and puberty are essential periods of postnatal brain maturation and are characterized by heightened susceptibility to mental disorders (Schneider, 2013). Specifically, changes in puberty onset are associated with increased risk for depression, anxiety (Stice et al., 2001; Kaltiala-Heino et al., 2003) and substance use (Hummel et al., 2013).

While essential for the maturation of adult social and cognitive skills (Casey et al., 2008), social relationships during adolescence are also implicated in the etiology of neuropsychiatric and neurodevelopmental disorders (Hankin et al., 1998). Social behavior is sexually dimorphic in rodents (Vanderschuren et al., 1997) and is, at least in part, controlled by the eCB system (Wei et al., 2017; Manduca et al., 2016; Manduca et al., 2015). Rather unsurprisingly, exposure to cannabinoid agonists during adolescence alters social behavior in adult rats (Schoch et al., 2018; Trezza and Vanderschuren, 2009; Trezza and Vanderschuren, 2008; Schneider et al., 2008).

The eCB system is differentially regulated according to sex (Cooper and Craft, 2018). Hormonal regulation affects eCB activity and sexual differences are apparent in the effects of cannabinoids. Human studies suggest sex differences in cannabis use (Cuttler et al., 2016; Schepis et al., 2011; Stinson et al., 2006; Gavranidou and Rosner, 2003). In rodents, the effects of cannabis differ between males and females especially around puberty (Wiley et al., 2017; Silva et al., 2016; Marusich et al., 2015; Rubino and Parolaro, 2015; Rubino et al., 2008; Casey et al., 2008).

Although the consequences of chronic exposure to cannabinoids during the adolescent period have been intensely studied (Hoffman et al., 2003; Lupica et al., 2004; Pistis et al., 2004; Liu et al., 2010; Cass et al., 2014; Lovelace et al., 2015a; for review see Zlebnik and Cheer, 2016), the neuronal and behavioral consequences of cannabis initiation, i.e. the first exposure to the drug, are less clear. A single exposure to THC in-vivo ablates eCB-mediated synaptic plasticity (i.e. short and long-term depression, LTD) in the accumbens and hippocampus (Mato et al., 2004) but not hippocampal CA1 long-term potentiation (LTP) (Hoffman et al., 2007) or eCB-LTD at VTA GABA synapses (Friend et al., 2017). Additionally, acute canannabinoid exposure impaired LTP in the ventral subiculum-accumbens pathway (Abush and Akirav, 2012). Thus, it appears that the effects of a single cannabinoid exposure greatly depend on the brain area.

An important caveat is that most of the aforementioned studies used adolescent rats which range in age between 25 and 45 days-old and does not take into account the pubertal period, i.e., its onset or completion. During this interval, the different phases of adolescence, early, mid-and late adolescence, are comprised, and are common for males and females. However, mid-adolescence, when the physical markers of puberty typically appear, differs between sexes: females reach puberty around post-natal day (PND) 30 to 40 while puberty takes place in males later at aproximately PND 40 to 50 (Burke et al., 2017; Vetter-O’Hagen and Spear, 2012; Schneider, 2008). Thus, based on the developmental profile of the eCB system and the sensitivity of the pubertal period, we reasoned that two factors, onset of puberty and sex, may further complexify the situation reagarding the effects of acute exposure to exogenous cannabinoids. For the present study we therefore decided to focus on pubescent and adult rats of both sexes who were tested for social and cognitive behaviors as well as neuronal and synaptic parameters in pyramidal neurons of the PFC 24 h after a single in-vivo cannabinoid exposure (SCE).

## Material and Methods

### Animals

Wistar rats bred in our animal facility were weaned from the mother at postnatal day (PND) 21 and housed in groups of 5 individuals of the same sex with 12 h light/dark cycles and *ad libitum* access to food and water. All experiments were performed in accordance with the European Communities Council Directive (86/609/EEC) and the United States National Institutes of Health *Guide for the care and use of laboratory animals*. All behavioral and electrophysiological experiments were performed on pubescent and adult rats from both sexes. Take into account that male and female rats do not reach puberty at the same time (Thomazeau et al., 2014a; Schneider, 2013), experiments in pubsecent animals were performed when male rats were 42-55 days in age and female rats were 30-40 days in age. Male and female rats were considered adult at PND 90-120. As pubescent males and females differ in age, the term “age-matched” used in the text referes to rats belonging to the same period, i.e., puberty or adulthood. All animals were experimentally naïve and used only once.

### Drugs

The CB1/CB2 cannabinoid agonist WIN55,212-2 (WIN; 2mg/kg) was dissolved in 10% polyethylene glycol/10% Tween80/saline and injected subcutaneously (s.c.) 24 h before the behavioral and electrophysiological essays. Control animals (Sham group) received vehicle. Solutions were freshly prepared on the day of the experiment and were administered in a volume of 2mL/kg for rats weighing <150 g and 1 mL/kg for adult rats. The 2 mg/kg dose chosen for single exposure is within the 1.2 to 3 mg/kg range that reliably causes behavioral and neuronal effects when given chronically (Wegener and Koch, 2009; Tagliaferro et al., 2006).

### Behavioral paradigms

The experiments were performed in a sound attenuated chamber under dim light conditions (15-25 lux). Animals were handled 2 consecutive days before starting the behavioral tests and adapted to the room laboratory conditions 1 hour before the tests. They were tested in a 45 x 45 cm arena with ±2 cm of wood shavings covering the floor. Drug treatments were counterbalanced by cage (mates were allocated to different treatment groups). Behavioral procedures were performed between 10:00 am and 3:00 pm. All sessions were recorded using a video camera using the Ethovision XT 13.0 video tracking software (Noldus, The Netherlands) and analysed by a trained observer who was unaware of treatment condition.

### Social behavior in pubescent and adult rats

The social behavior test was performed as previously published (Manduca et al., 2015). The animals of each pair were equally treated (WIN or vehicle), did not differ more than 10 g in body weight and were sex and age mates but not cage mates. Pubescent or adult rats of both sexes were individually habituated to the test cage daily for either 10 (pubescent) or 5 min (adult) 2 days prior to testing. At the end of the second day of habituation (24 h before the test), the rats received the treatment. To enhance their social motivation and thus facilitate the expression of social behaviors, pubescent and adult animals were socially isolated before testing for 3.5 and 24 h, respectively (Niesink and Van Ree, 1989). The test consisted of placing two equally treated rats into the test cage for either 15 min (pubescent) or 10 min (adult).

In pubescent rats, we scored: 1/ Social behavior related to play: pouncing (one animal is soliciting the other to play by attempting to nose or rub the nape of its neck) and pinning (one animal lying with its dorsal surface on the floor with the other animal standing over it). This is the most characteristic posture in social play in rats; it occurs when one animal is solicited to play by its test partner and rotates to its dorsal surface (Panksepp and Beatty, 1980; Trezza et al., 2010) and 2/ Social behavior unrelated to play (assessed as a measure of general social interest): sniffing (when the rat sniff, licking, or grooms any part of the body of the test partner).

In adult rats we scored: 1/ Play-related behaviors: pouncing, pinning and boxing and 2/ Social behaviors unrelated to play: sniffing, social grooming (the rat licks and chews the fur of the conspecific, while placing its forepaws on the back or the neck of the other rat), following/chasing (walking or running in the direction of the partner which stays where it is or moves away), crawling under/over (one animal crawls underneath or over the partner’s body, crossing it transversely from one side to the other), kicking (the rat kicks backwards at the conspecific with one or both hind paws).

The parameters were analysed grouped and considered as *total social exploration*, calculated as the sum of social behaviors. Aggressive behavior was also scored but not considered in the calculation of *total social exploration*.

### Novel object recognition test

The test comprised two phases: training (acquisition trial) and test. Each session lasted 5 minutes. During the acquisition trial, the rat was placed into the arena containing two identical sample objects (A1 and A2) placed near the two corners at either end of one side of the arena (8 cm from each adjacent wall). Thirty minutes later, the rat returned to the apparatus containing two objects, one of them was a copy to the object used in the acquisition trial (A3), and the other one was novel (B). The objects in the test were placed in the same positions as during the acquisition trial. The positions of the objects in the test and the objects used as novel or familiar were counterbalanced between the animals. Exploration was scored when the animal was observed sniffing or touching the object with the nose and/or forepaws. Sitting on objects was not considered to indicate exploratory behaviour. The apparatus and the objects were cleaned thoroughly with 50% ethanol between trials to ensure the absence of olfactory cues. The discrimination ratio was calculated as follow: time spent by each animal exploring the novel object divided by the total time spent exploring both objects. Discrimination ratio higher than 0.5 indicates preferable object recognition memory. Number of rearing and grooming were registered during the acquisition trial.

### Slice preparation

Twenty-four hours after WIN or vehicle administration, rats were anesthetized with isoflurane and decapitated according to institutional regulations. The brain was sliced (300 µm) in the coronal plane with a vibratome (Integraslice, Campden Instruments, Loughborough, UK) in a sucrose-based solution at 4°C (values in mM: 87 NaCl, 75 sucrose, 25 glucose, 5 KCl, 21 MgCl_2_, 0.5 CaCl_2_, and 1.25 NaH_2_PO_4_). Slices were allowed to recover for 60 min at ±32°C in a low calcium artificial cerebrospinal fluid (aCSF) (in mM: 126 NaCl, 2.5 KCl, 2.4 MgCl_2_, 1.2 CaCl_2_, 18 NaHCO_3_, 1.2 NaH_2_PO_4_, and 11 glucose, equilibrated with 95% O_2_/5% CO_2_. Slices were maintained at room temperature until recording.

### Electrophysiology

Whole-cell patch-clamp and extra-cellular field recordings were made from layer 5 pyramidal cells of the prelimbic cortex (mPFC) (Martin et al., 2016b; Kasanetz et al., 2013). For recording, slices were superfused (1.5–2 mL/min) with aCSF containing picrotoxin (100 μM) to block GABA_A_ receptors. All experiments were performed at 32±2°C. To evoke synaptic currents, 100–200 μs stimuli were delivered at 0.1 Hz through an aCSF-filled glass electrode positioned dorsal to the recording electrode in layer 5. Patch-clamp recordings were performed with a potassium gluconate based intracellular solution (values mM: 143 potassium gluconate, 3 NaCl, 1 MgCl_2_, 0.3 CaCl_2_, 1 EGTA, 0.2 cAMP, 0.3 NaGTP, 2 NaATP, 10 HEPES, pH 7.25, osmolarity 290–300 mol/L). Patch pipettes had a resistance between 3 and 5 MΩ. In all experiments cells were clamped at −70 mV (without junction potential correction). During recordings holding currents, series and input resistances and the membrane time constant (τ) were monitored. If the series resistance exceeded 25 MΩ or varied by >20% during the experiment the recording was rejected.

Current-voltage (*I–V*) curves were made by a series of hyperpolarizing to depolarizing current steps immediately after breaking into the cell. Membrane resistance was estimated from the *I–V* curve around resting membrane potential (Thomazeau et al., 2014).

For extracellular field experiments, the recording pipette was filled with aCSF. The glutamatergic nature of the field excitatory postsynaptic potential (fEPSP) was systematically confirmed at the end of the experiments using the ionotropic glutamate receptor antagonist 6-cyano-7-nitroquinoxaline-2,3-dione (CNQX, 20 μM), that specifically blocked the synaptic component without altering the non-synaptic component (data not shown). Example EPSPs and fEPSPs are single sweeps from the indicated time points, for clarity the stimulation artefact was removed from the fEPSP.

### Data analysis

The magnitude of plasticity was calculated 35–40 min after and compared to the average of baseline response. sEPSCs were analyzed with Axograph X (Axograph). Statistical analysis of data was performed with Prism 6 (GraphPad Software) using tests indicated in the main text after outlier subtraction (Grubb’s test). Grafical values are given as mean±SEM and table values are given as median and interquartiles ranges. Statistical significance was set at *p*<0.05 (two-tailed).

## Results

### Single exposure to WIN alters social behavior in a sex-and age-dependent manner

We compared distinct behavioral elements related to the social repertoire of rodents in male and female rats at different ages (puberty and adulthood) previously exposed to a single dose (2 mg/kg) of the synthetic cannabinoid agonist WIN55,212-2 (WIN). In contrast with previous studies where animals were tested shortly after after WIN administration, i.e. 30 min after 0.1–1 mg/kg (Trezza and Vanderschuren, 2008a), 0.3 mg/kg (Trezza and Vanderschuren, 2008b) and 1.2 mg/kg (Schneider et al., 2008), the behavioral and synaptic tests were performed 24 h after WIN administration to take advantage of WIN’s short half-life of terminal elimination (5h) (Valiveti et al., 2004).

At puberty, male rats exhibited normal social play behavior 24 h after SCE: the number of pouncing (Fig. 1A: *U*=44, p=0.696, Mann-Whitney *U*-test) and pinning (Fig. 1C: *U*=42, p=0.588, Mann-Whitney *U*-test) behaviors were unaltered. Accordingly, the total time spent exploring the partner during the test was unaffected (Fig. 1E: *U*=42, p=0.602, Mann-Whitney *U*-test). In contrast, female rats at this same age showed significant reduction on parameters related to play behavior evidenced by a marked reduction in the number of play solicitations, i.e., pouncing (Fig. 1B: *U*=13.5, p=0.008, Mann-Whitney *U*-test) and play responses, i.e., pinning (Fog. 1D: *U*=9, p=0.001, Mann-Whitney *U*-test) observed 24 h after WIN administration. On the other hand, the total time spent exploring the social partner was comparable to that of the Sham group (Fig. 1F: *U*=27, p=0.156, Mann-Whitney *U*-test), indicating a specific impairment on social play behavior in pubescent females.

**Figure 1.**
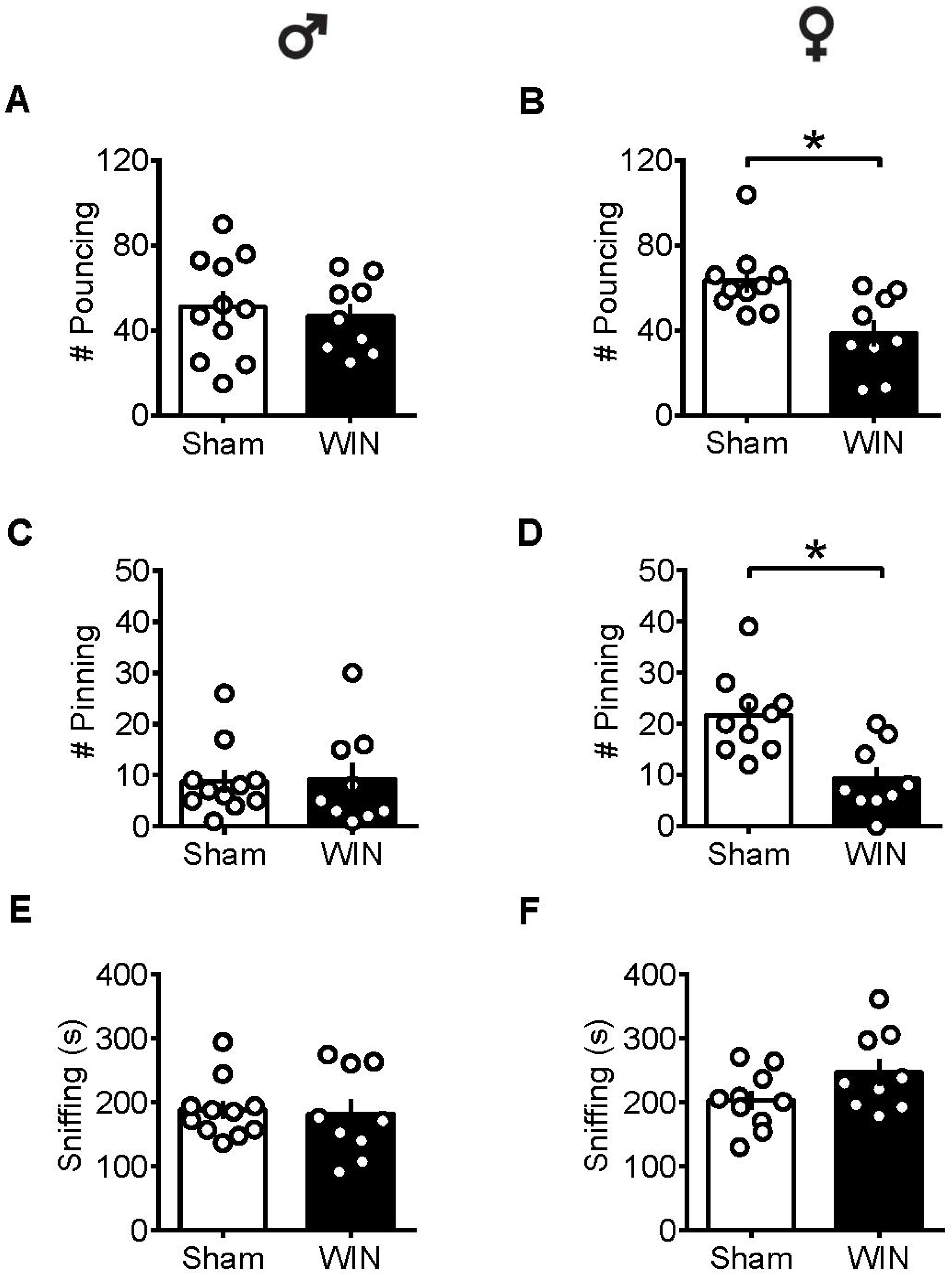
Sex-specific alteration of play behavior in pubescent rats 24 h after a single in-vivo cannabinoid exposure. 24 h following a single exposure to WIN55,212-2 (WIN, 2 mg/kg, s.c.), pouncing was normal in male pubescent rats (A) in contrast to female littermates whom displayed a marked reduction in the number of pouncing compared to Sham animals. 24 h following WIN exposure, pinning was similar to that of Sham animals in males but was largely reduced in female littermates (D). WIN-exposed rats of both sexes (E, male and F, female) spent similar time sniffing the congener compared to their respective Sham groups. Data represent mean ± SEM. Scatter dot plot represents a pair of animals. *p<0.05, Mann-Whitney *U*-test. ♂ males; ♀ females.

In contrast to pubescent rats, both male and female adult rats showed reduced social interest 24 h after SCE. Adult male rats administered WIN presented reduced general social exploration (Fig. 2A: *U*=7, p=0.003, Mann-Whitney *U*-test) as well as reduced sniffing exploration (Fig. 2C: *U*=3.5, p<0.001, Mann-Whitney *U*-test) compared to the Sham group. Similarly, adult cannabinoid-exposed females had less social contact (Fig. 2B: *U*=14.5, p=0.007, Mann-Whitney *U*-test) and sniffing events (Fig. 2D: *U*=15.5, p=0.010, Mann-Whitney *U*-test) with congeners. In addition, SCE did not elicit aggressive behavior in any of the tested groups (data not shown).

**Figure 2.**
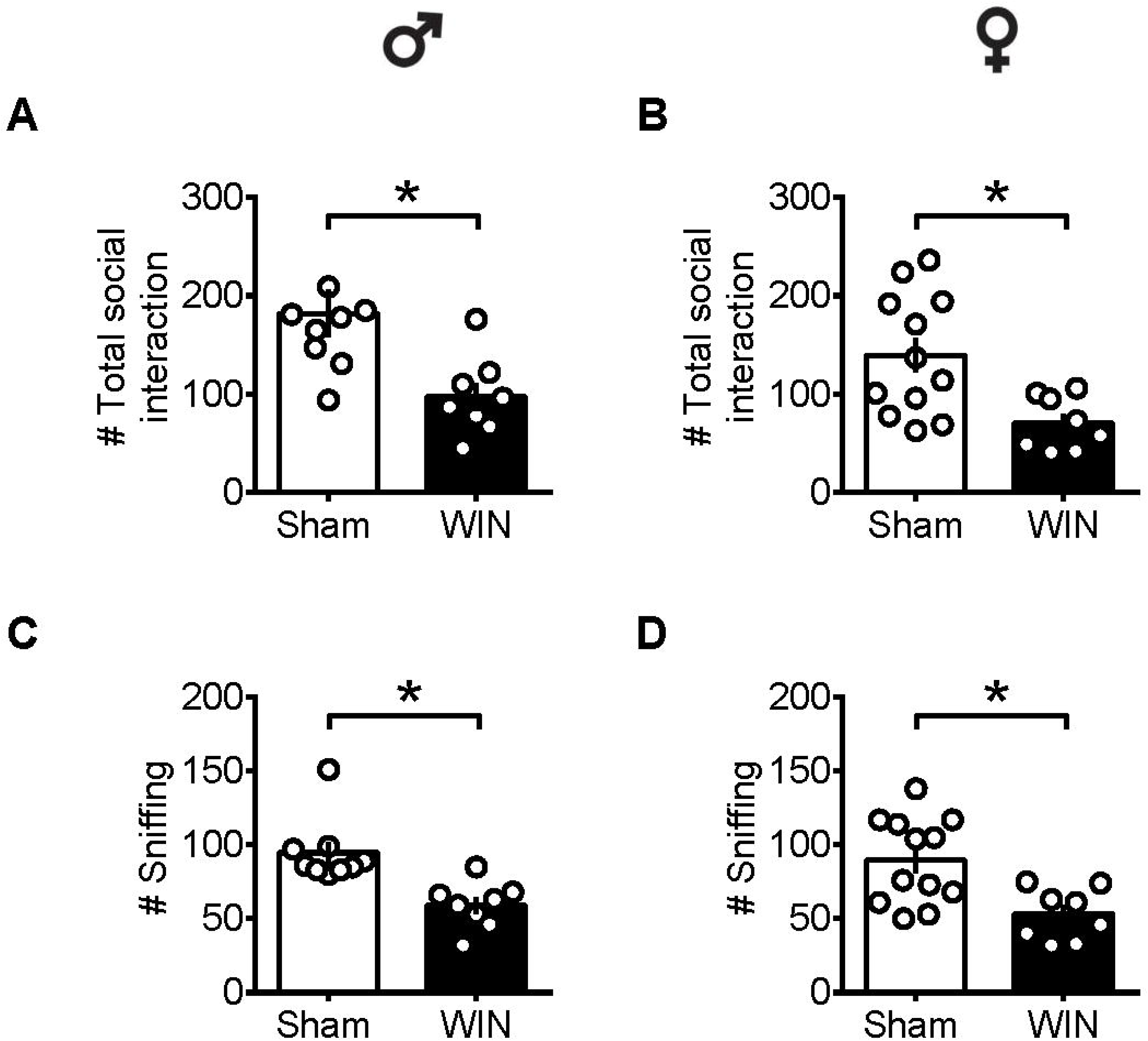
Social interactions are diminished in adult rats of both sex 24 h after a single in-vivo cannabinoid exposure. Adult male (A) and female (B) rats had less social contacts with their congeners 24 h following a single exposure to WIN. Similarly, sniffing was reduced in both adult male (C) and female (D) rats 24 h following a single exposure WIN, compared to control animals. Data represent mean ± SEM. Scatter dot plot represents a pair of animals. *p<0.05, Mann-Whitney *U*-test. ♂ males; ♀ females.

Together, these data show that during puberty, SCE is sufficient to alter social behavior in a sex-specific manner: play behavior was specifically reduced in females while males were spared. In adults, SCE caused a general impairment in sociability, exhibited by a reduced number of events related to general exploration and sniffing in both male and female rats.

Importantly, we showed that the low socialization observed in pubescent female rats and in adult rats of both sexes was unlikely due to an impaired exploration since behavioral parameters unrelated to cognition but linked to general exploration and emotionality, as rearing and grooming occurrences, were unchanged 24 h after SCE (Table 1).

**Table 1.** Statistical report for general exploration parameters (rearing and grooming) in pubescent and adult rats from both sexes 24 h after a single in-vivo exposure to WIN (2 mg/kg, s.c.). The number of rearing and grooming was counted during the acquisition trial in the novel object recognition test. Quartiles, 25 and 75% percentiles; n, number of animals; Mann-Whitney *U*-test; ♂ males; ♀ females.

|  |  | Sham |  |  | WIN |  |  | *p* | *U* |
| --- | --- | --- | --- | --- | --- | --- | --- | --- | --- |
|  |  | Median | Quartiles | n | Median | Quartiles | n |  |  |
| Rearing | ♂ Pubescent | 37.50 | 32.75/46.00 | 10 | 38.00 | 31.00/43.50 | 9 | 0.826 | 42 |
|  | ♂ Adult | 35.50 | 25.25/44.00 | 8 | 36.00 | 16.50/40.00 | 9 | 0.524 | 29 |
|  | ♀ Pubescent | 35.00 | 25.00/40.50 | 9 | 37.00 | 30.50/45.00 | 13 | 0.502 | 48 |
|  | ♀ Adult | 22.00 | 19.50/30.50 | 5 | 28.00 | 23.00/44.00 | 7 | 0.162 | 8.5 |
| Grooming | ♂ Pubescent | 0.50 | 0/2.25 | 10 | 2.00 | 1.00/3.00 | 9 | 0.129 | 26.5 |
|  | ♂ Adult | 1.00 | 0.25/1.75 | 8 | 1 | 1.00/3.00 | 9 | 0.395 | 26.5 |
|  | ♀ Pubescent | 1.00 | 0/2.50 | 9 | 1 | 1.00/5.00 | 13 | 0.732 | 53 |
|  | ♀ Adult | 2.00 | 0.50/4.50 | 5 | 3.00 | 0/5.00 | 7 | 0.977 | 17 |

### Intact memory recognition in pubescent and adult rats of both sexes after single cannabinoid exposure

In humans (Walsh et al., 2017) and rodents (Wegener et al., 2008; Han et al., 2012; Galanopoulos et al., 2014), cannabinoids rapidly impair recent memory. Social behavior requires emotional control and cognitive abilities (Trezza et al., 2014). Thus, we used the novel object recognition test to evaluate the consequences of SEC on rats of our sex and age groups. 24 h after SCE, pubescent male (Fig. 3A: *U*=17, p=0.999, Mann-Whitney *U*-test) and female (Fig. 3B: *U*=52, p=0.682, Mann-Whitney *U*-test) rats presented normal short-term memory. Furthermore, discrimination indexes were similar in both adult male and female Sham-and WIN-treated rats (Fig. 3C: male, *U*=29.5, p=0.557; Fig. 3D: female, *U*=15, p=0.755; Mann-Whitney *U*-test). Importantly, the total time spent exploring the objects during the acquisition trial was not altered in any of the tested groups (Pubescent Males: Sham *vs.* WIN, *U*=31, p=0.277; Pubescent Females: Sham *vs*. WIN, *U*=42, p=0.292; Adult Males: Sham *vs.* WIN, *U*=31, p=0.673; Adult Females: Sham *vs*. WIN, *U*=5, p=0.082; Mann-Whitney *U*-test; data not shown).

**Figure 3.**
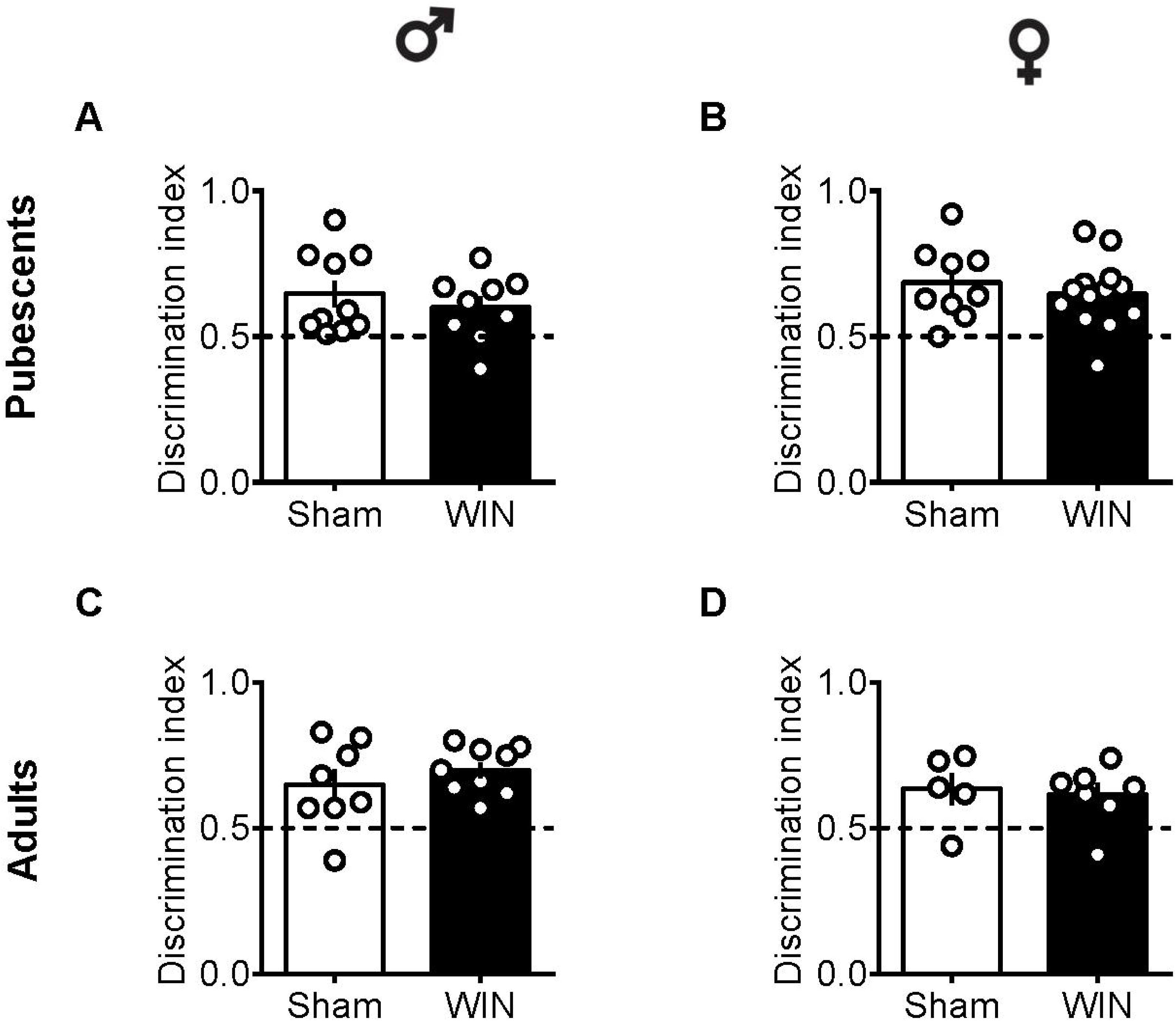
Intact memory discrimination in the novel object recognition test 24 h after single in-vivo cannabinoid exposure in both pubescent and adult male and female rats. Discrimination ratio between the novel and familiar objects were similar in male (A) and female (B) WIN-treated pubescent rats compared to their respective Sham groups. Similarly, no differences were observed in discrimination ratio in male (C) and female (D) adult rats treated with WIN. Data represent mean ± SEM. Scatter dot plot represents one animal. Mann-Whitney *U*-test. ♂ males; ♀ females.

### Single in-vivo cannabinoid exposure leads to sex-specific ablation of prefrontal eCB plasticity

The central position of the PFC and eCB system in the regulation of social behavior and the important role of synaptic plasticity in this structure in mediating experience-dependent adaptations are well-documented (for review see Araque et al., 2017). At the synaptic level, activity-dependent plasticity in the PFC – including eCB-mediated long-term depression (LTD) and NMDAR-mediated long-term potentiation (LTP) – is a common target in animal models of neuropsychiatric diseases (Scheyer AF et al., 2017). We compared the LTD mediated by the eCB system (eCB-LTD) in the PFC between Sham-and WIN-treated rats of both sexes at different ages, specifically pubescence and adulthood.

Low-frequency stimulation of layer V PFC synapses induced comparable LTD in both control and cannabinoid-exposed pubescent male rats (Fig. 4A: Sham: t_(6)_=5.596, p=0.001; WIN: t_(4)_=3.190, p=0.033; Paired *t*-test). Similar results were observed in adult male with or without prior in-vivo cannabinoid exposure (Fig. 4B: Sham, t_(6)_=3.116, p=0.020; WIN, t_(6)_=2.787, p=0.031; Paired *t*-test). In contrast to what we observed in male mouse hippocampus and accumbens in a previous study (Mato et al., 2004), it appears that in the male rat PFC, eCB-LTD is not affected 24 h after in-vivo cannabinoid administration. Strikingly, eCB-LTD was ablated in PFC slices obtained from female rats in both age groups. Figure 4C shows the lack of LTD in PFC slices from cannabinoid-treated pubescent (Sham, t_(4)_=5.021, p=0.007; WIN, t_(4)_=1.129, p=0.322; Paired *t*-test) and adult female rats (Fig. 4D: Sham, t_(4)_=2.979, p=0.040; WIN, t_(7)_=1.003, p=0.349; Paired *t*-test).

**Figure 4.**
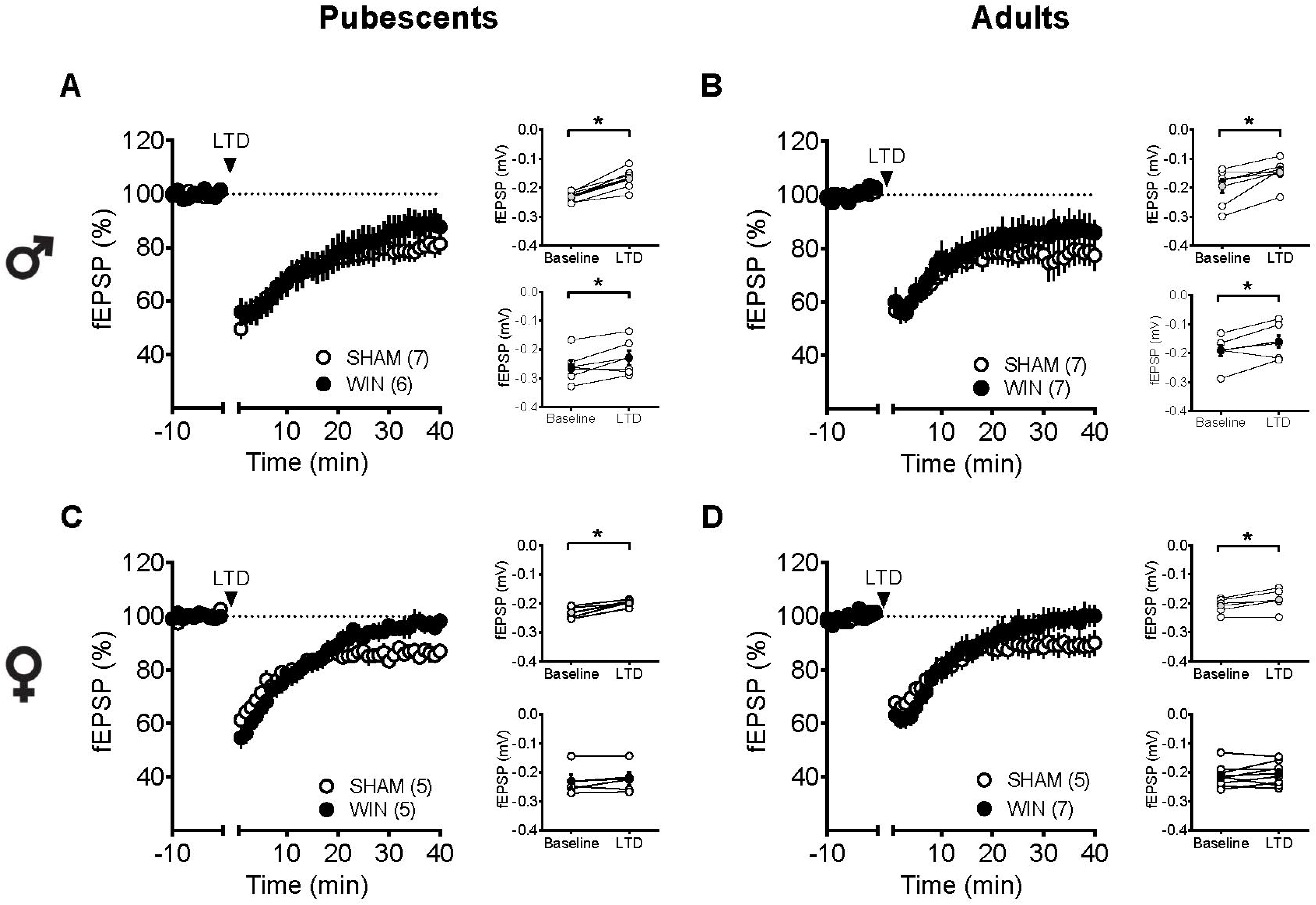
Sex-specific effects of a single in-vivo cannabinoid exposure on PFC eCB-LTD. Average time-courses of mean fEPSPs showing that low-frequency stimulation (indicated by arrow) induced LTD at mPFC synapses in both Sham-(white circles, n=7) and WIN-(black circles, n=6) exposed pubescent males (A). Similarly, LTD was identical in Sham-(white circles, n=7) and WIN-(black circles, n=7) exposed adult males (B). In contrast, LTD was ablated in mPFC slices obtained from both pubescent (C, Sham, white circles, n=5; WIN, black circles, n=5) and adult (D, Sham, white circles, n=5; WIN, black circles, n=7) females 24 h after a single exposure to WIN. Adjacent to the time-course figures individual experiments (white circles) and group average (Sham, gray circles; WIN, balck circles) before (baseline) and after (35-40 min) LTD induction are showed. LTD is present in WIN-treated male rats at both ages: pubescent (A, on the rigth) and adulthood (B, on the rigth). In contrast, LTD was absent in both pubescent (C, on the right) and adult (D, on the right) females previously treated with WIN. Error bars indicate SEM, n= individual rats, *p<0.05, Paired *t*-test. ♂ males; ♀ females.

### Age-and sex-dependent ablation of LTP after in-vivo single expsoure to cannabinoid

Considering that the extensive repertoire of synaptic plasticity expressed by medial PFC synapses is sensitive to various regimen of exposure to drugs of abuse (Kassanetz et al. 2010; Cannady et al., 2017, Renard et al., 2016; Lovelace et al., 2015; van Huijstee and Mansvelder, 2014) we assessed a second type of plasticity in the PFC which is frequently related to endophenotypes of neuropsychiatric disorders (Labouesse et al. 2016; Manduca et al., 2017; Neuhofer et al., 2015; Iafrati et al., 2016; Thomazeau et al., 2014), the NMDAR-dependent LTP (NMDAR-LTP). NMDAR-LTP was ablated in adult male rats while pubescent males were spared. Figures 5A-B show comparable LTP between Sham and cannabinoid-treated pubescent male rats (Fig. 5A: Sham, t_(6)_=9.676, p<0.001; WIN, t_(7)_=3.677, p=0.007; Paired *t*-test), but not in adult male rats (Fig. 5B: Sham, t_(8)_=5.560, p<0.001; WIN, t_(6)_=2.062, p=0.084; Paired *t*-test). In contrast, in both age groups, NMDAR-LTP was comparable in Sham and cannabinoid-treated female rats: both pubescent (Fig. 5C: Sham, t_(6)_=8.424, p<0.001; WIN, t_(6)_=3.369, p=0.015; Paired *t*-test) and adult rats (Fig. 5D: Sham, t_(4)_=4.349, p=0.012; WIN, t_(7)_=3.133, p=0.016; Paired *t*-test) had normal NMDAR-LTP 24 h following in-vivo cannabinoid exposure.

**Figure 5.**
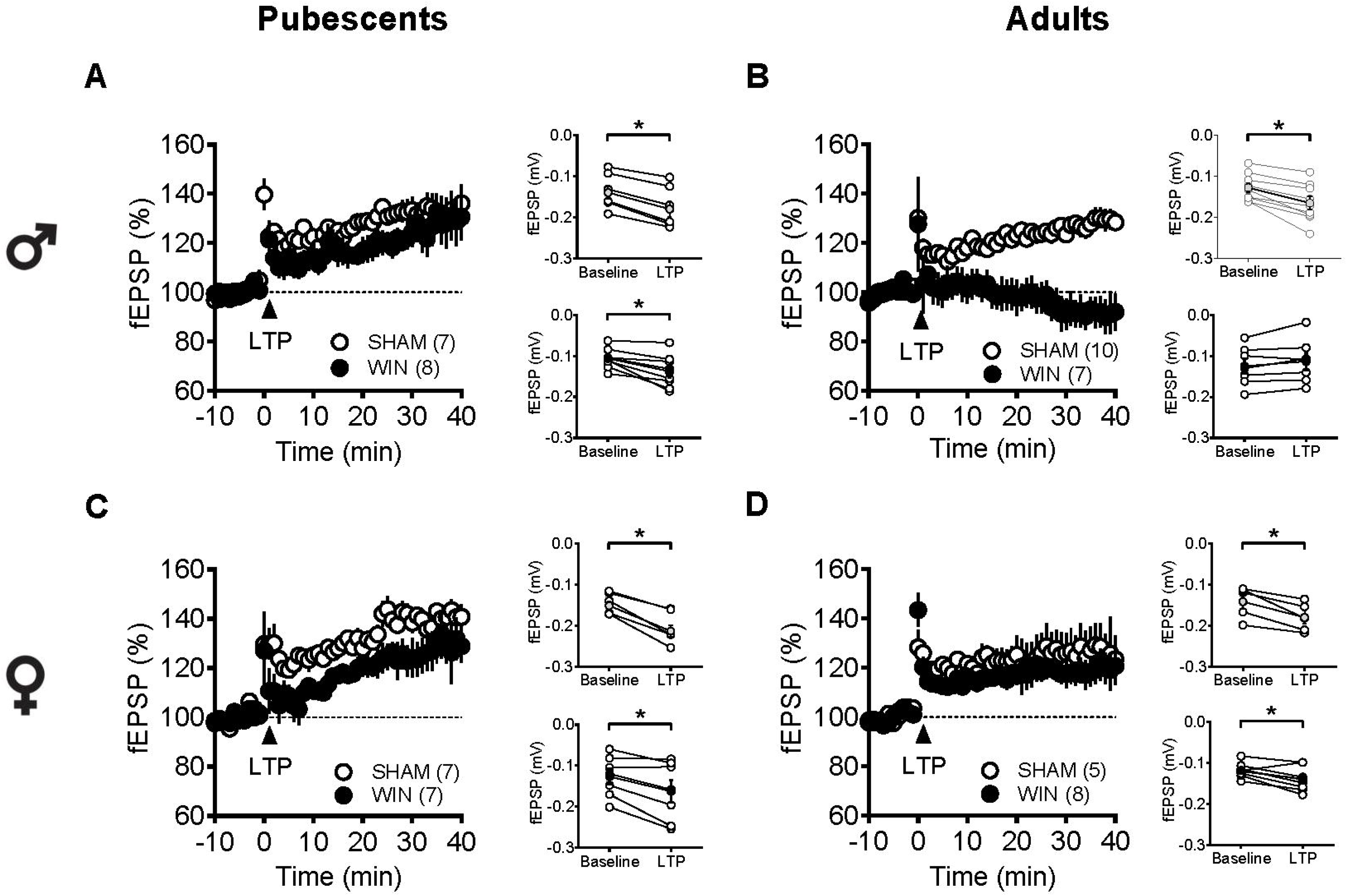
Age-and sex-dependent ablation of LTP in the rat PFC 24 h after in-vivo cannabinoid exposure. Average time-courses of mean fEPSPs showing that theta-burst stimulation (indicated arrow) induced a LTP at mPFC synpases in both Sham-(white circles, n=7) and WIN-(black circles, n=8) exposed pubescent males (A) but not WIN-treated adults (B, Sham: white circles, n=10; WIN: black circles, n=7). In contrast, LTP was present in mPFC slices obtained from both pubescent (C, Sham: white circles, n=7; WIN: black circles, n=7) and adult (D, Sham: white circles, n=5; WIN: black circles, n=8) WIN-treated females. Adjacent to the time-course figures are showed individual experiments (white circles) and group average (Sham, gray circles; WIN, black circles) before (baseline) and after (35-40 min) LTP induction showing that, in males, LTP is present in pubescent (A, on the right) but not in adults (B, on the right). In contrast, LTP was present in both pubescent (C, on the right) and adult (D, on the right) females previously treated with WIN. Error bars indicate SEM, n= individual rats, *p<0.05, Paired *t*-test. ♂ males; ♀ females.

### Single in-vivo exposure to WIN causes age-and sex-specific modifications in intrinsic pyramidal neuron properties

Independent of sex, all recorded PFC neurons in pubescent rats showed similar membrane reaction profiles in response to a series of somatic current steps 24 h after SCE (Fig. 6A: Male, F_(interaction10,440)_=1.551, p=0.118; Fig. 6B: Female, F_(interaction10,270)_=0.499, p=0.889; two-way repeated-measures ANOVA). The resting membrane potential (Fig. 6C: Male, *U=* 230, p=0.627; Fig. 6D: Female, *U*=99.5, p=0.854; Mann-Whitney *U*-test) as well as the rheobase (Fig. 6E: Male, *U*=194.5, p=0.198; Fig. 6F: Female, *U*=68, p=0.115; Mann-Whitney *U*-test) were comparable between Sham and WIN-treated pubescent rats from both sexes. Also, no changes in excitability were observed since the number of actions potentials in response to somatic currents steps were comparable in both control and WIN-treated pubescent rats of both sexes (Fig 6G: Male, F_(interaction 12,492)_=1.189, p=0.287; Fig. 6H: Female, F_(interaction_ 12,324)=3.624, p<0.001 and F_(treatment 1,27)_=0.389, p=0.537; two-way repeated measures ANOVA).

**Figure 6.**
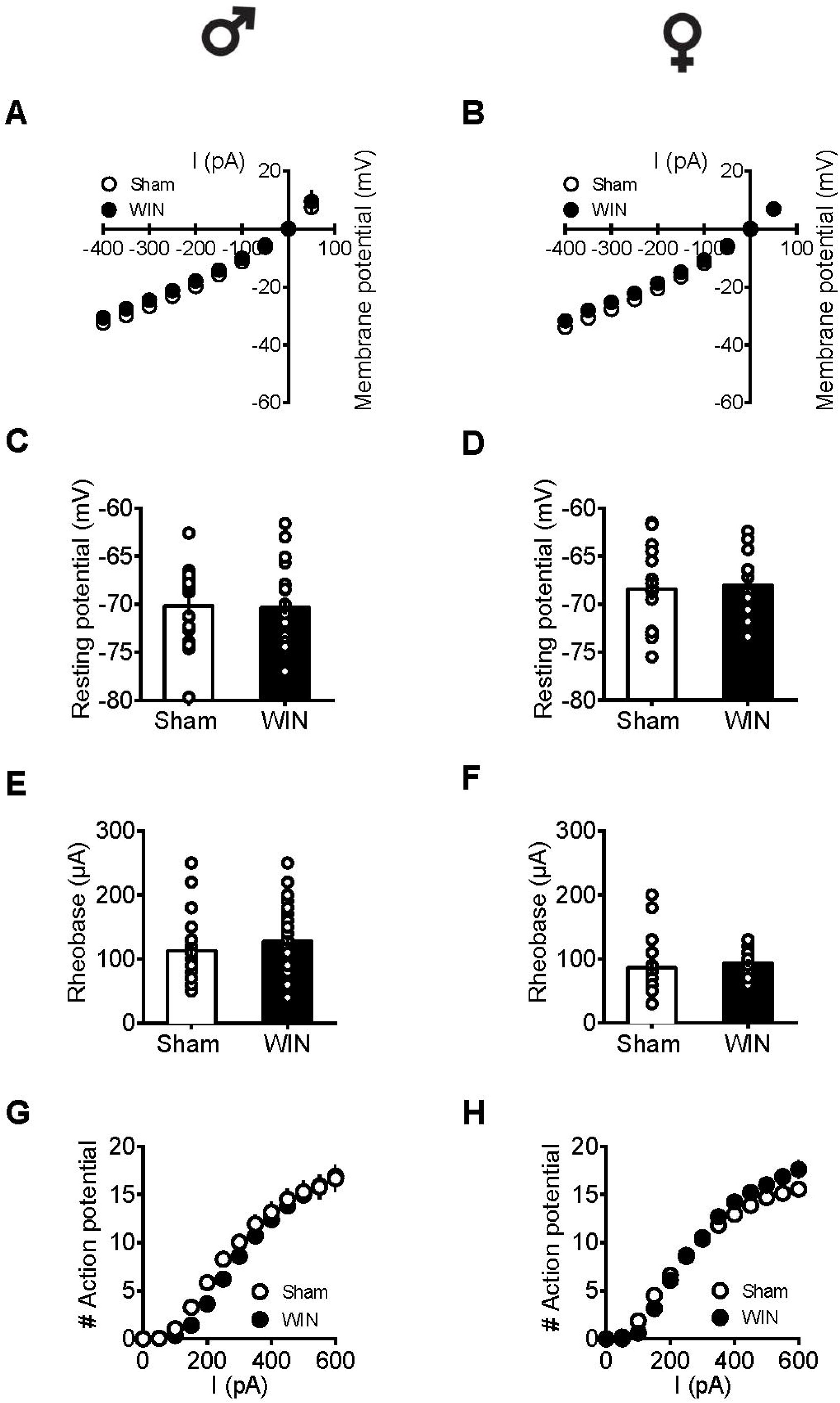
Intrinsic properties of PFC pyramidal neurons are not altered by single in-vivo exposure to cannabinoid in pubescent rats. Current–voltage plot from visually identified pyramidal neurons recorded from pubescent rats showing similar cell voltage in response to current steps between Sham and WIN of both male (A) and female (B) groups. No change was observed in the resting membrane potential 24 h after WIN treatment in both male (C) and female (D) pubescent rats. Quantification of neuronal spiking properties indicated no change in the rheobase of either males (E) or females (F) 24 h after single WIN. The number of evoked action potentials in response to increasing depolarizing current steps was similar in Sham and WIN-treated male (G) and female (H) pubescent rats. Males: Sham, n=15 cells/6 rats; WIN, n=28 cells/10 rats. Females: Sham, n=17 cells/7 rats; WIN, n=13 cells/7 rats. Scatter dot plot represents one cell. Data represent mean ± SEM. ♂ males; ♀ females.

In adult rats however, sex-specific modifications of the excitability of pyramidal neurons sampled from females were observed following a single in-vivo cannabinoid exposure. Intrinsinc properties of layer V PFC pyramidal neurons were comparable in control and WIN-treated male rats (I/V curve Fig. 7A: F_(interaction 9,225)_=1.907, p=0.052, two-way repeated measures ANOVA) resting membrane potentials (Fig. 7C: *U*=79, p=0.614, Mann-Whitney *U*-test; rheobase Fig. 7E: *U*=79, p=0.614, Mann-Whitney *U*-test) and the number of action potentials in response to increasing depolarizing current (Fig. 7G: F_(interaction 10,250)_=1.417, p=0.173, two-way repeated measures ANOVA). In striking contrast, a single in-vivo cannabinoid exposure increased the excitability of PFC pyramidal neurons of adult females. Thus, we observed an alteration of the membrane reaction profile in response to a series of somatic current steps (Fig. 7B: F_(interaction 9,369)_=3.480, p<0.001 and F_(treatment 1,41)_=5.576, p=0.023, two-way repeated measures ANOVA) and a marked reduction of the rheobase (Fig. 7F: *U*=137.5, p=0.023, Mann-Whitney *U*-test) accompanying an increased number of action potential in response to increasing depolarizing current (Fig. 7G: F_(interaction 10,410)_=3.038, p=0.001 and F_(treatment 1,41)_=8.041, p=0.007, two-way repeated measures ANOVA). The resting membrane potentials were similar to that of control female rats (Fig. 7D, *U*=166.5, p=0.124, Mann-Whitney *U*-test). Taken together, these data suggest an overall increase in the excitability of PFC pyramidal neurons in adult females 24H after SCE.

**Figure 7.**
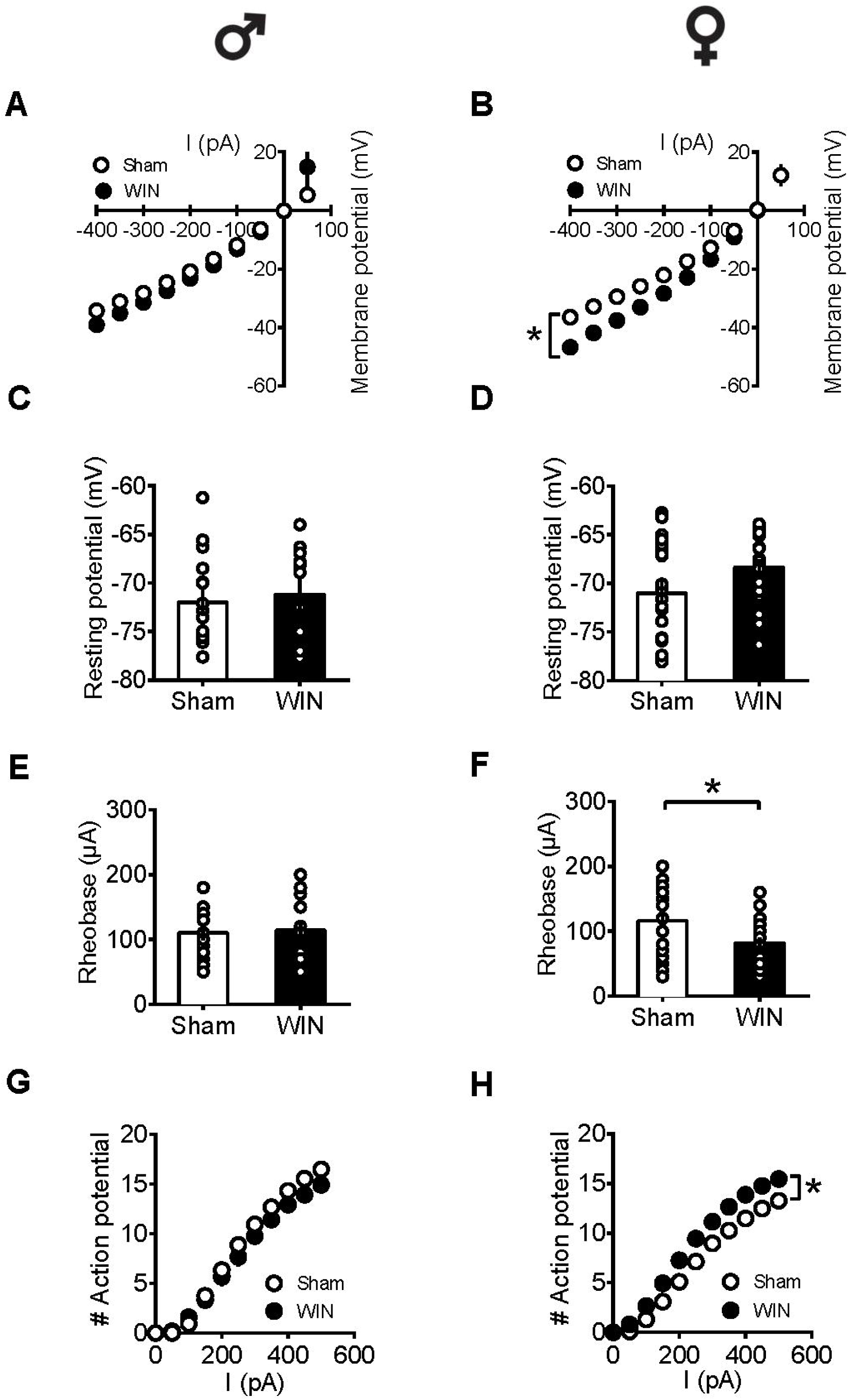
Sex-specific alteration of pyramidal neurons’ intrinsic properties in adult rats 24 h following single in-vivo cannabinoid administration. Current–voltage plot from visually identified pyramidal neurons recorded from adult rats showing no difference in cell voltage in response to current steps between Sham and WIN groups (green symbols) of adult male rats (A). In contrast, membrane potentials were altered in adult WIN-treated females compared to control group (B). The resting membrane potentials were similar to that of control in adult males (C) and females (D) 24 h following single WIN exposure. Quantification of neuronal spiking properties showed no change in the rheobase of males (E), but a marked reduction in the female WIN-treated group (F). The number of evoked action potentials in response to increasing depolarizing current step was similar in Sham and WIN-treated males (G). In contrast, females showed a higher number of action potentials 24 h after WIN treatment (H). Male: Sham, n=16 cells/6 rats; WIN, n= 12 cells/5 rats; Female: Sham, n=16 cells/6 rats; WIN, n=20 cells/7 rats. Scatter dot plot represents one cell. Data represent mean ± SEM. *p<0.05, Mann-Whitney test (B), Bonferroni’s multiple comparisons test (F). ♂ males; ♀ females.

## Discussion

We found that 24 h after a single in-vivo exposure to a cannabinoid, the behavioral, neuronal and synaptic consequences differ depending on the sex and age of the rat. The current data indicate a heightened sensitivity of females, especially during pubescence. Specifically, social behavior and eCB-mediated LTD showed strong deficits in exposed pubescent females while age-matched male littermates were spared. During adulthood, although reduced social interactions were observed in both sexes, eCB-mediated synaptic plasticity was ablated specifically in females and NMDAR-dependent LTP in males.

Stimulation of CB1R acutely modulates social play in adolescent rats (Trezza and Vanderschuren, 2008a). We showed that a single exposure to the synthetic cannabinoid WIN (2 mg/kg), at a dose reported to acutely decrease social interactions in male rats (Schneider et al., 2008; Trezza and Vanderschuren, 2008a; Trezza and Vanderschuren, 2008b) has sex-specifc effects as long as 24 h after in-vivo exposure. In the pubescent group, cannabinoid-treated females exhibited less social play behavior but normal social investigation, while the sociability of male littermates exposed to WIN was indistinguishable from that of sham rats. It is important to mention that pubescent female sham rats presented augmented number of pinnings when compared to age-matched sham males (*U*= 11, p=0.001, Mann-Whitney test, data not shown). However, data from the literature show that adolescent males have higher levels of play behavior than age-matched females (Argue and McCarthy, 2015; Burke et al., 2017). Our finding may be explained by the difference in the age range in which males and females were herein tested. Our objective was to verify the effect of acute cannabinoid exposure in pubescent rats regardeless of the onset of the adolescent period. Thus, considering that in our conditions the play behavior of pubescent females was reduced 24h after SCE to the same levels of those observed in control males, we may infer that SCE induced a masculinization of female social play behavior. Interestingly, a recent study showed that the activation of both CB1 and CB2 receptors (as that observed following exposure to WIN) is implicated in the masculinization of play behavior of pre-pubertal female rats (Argue et al., 2017), reinforcing the idea of sex-dependet modulation of social behaviors that arises early in life. Taken together these data confirm and extend those of Craft and collaborators (2013) who showed that females are more affected by exogenous cannabinoids during pubescence then males.

Interactions with age-matched congeners during adolescence are crucial for the development of social competence at adulthood (Douglas et al., 2004; Vanderschuren and Trezza, 2014) and modification of the rat adolescent social activity alters neurobehavioral parameters related to pain processing, anxiety, depression and substance abuse (reviewed from Burke et al., 2017). Thus, future experiments are necessary to determine if the deficits caused by SCE are long-lasting. Available data do not favor this scenario (Mato et al. 2004). Although sex differences on cannabinoids’ effects on cognition have been reported (Wiley et al., 2017; Silva et al., 2016; Marusich et al., 2015; Rubino and Parolaro, 2015; Rubino et al., 2008; Marco et al., 2006), in the present experiments neither locomotion nor novel object recognition memory were affected in either sex, in favor of the idea that the deficits are not generalized but rather selective to the social behavior.

Gonadal steroids hormones seem to be involved in the sexual differentiation of cannabinoid sensitivity. Importantly, rat hormonal status (i.e., estrous cycle phase) has been reported to significantly influence sex differences for cannabinoid effects (revised from Cooper and Craft, 2018). Indeed, sex differences are not entirely consistent across studies regarding differences in CB1R mRNA or binding affinity and eCB content (Weed et al., 2016; Castelli et al., 2014; Riebe et al., 2010; Reich et al., 2009), supporting the important role of hormonal status in these differences.

In contrast to the pubescent groups, a unique exposure to WIN triggered a different response in adults, since both sexes exhibited perturbed social behaviors. Adolescent rodents are more sensitive to cannabis than adults (Renard et al., 2016a). Surprisingly, here we showed that 24 h after SCE, pubescent males did not display behavioral or synaptic changes, while adult rats did. As cannabinoid doses, administration route, post-administration intervals and rat strains are not consistent among studies, methodological details may help explaining this discrepancy. In addition, we cannot rule out a potential protective effect of gonadal hormones in pubescent rats, since testosterone protects gonadectomized males against THC dependence (Marusich et al., 2015b). Thus, considering that this gonadal hormone reaches its peak during the pubertal period (Pignatelli et al., 2006), we can speculate that testosterone “protected” pubescent males from the residual deleterius effect of cannabinoids on social behavior and PFC synaptic plasticity.

Evidence shows that chronic cannabinoid exposure significantly impairs synaptic plasticity throughout the brain (Renard et al., 2016a; Araque et al., 2017), while the synaptic plasticity deficits resulting from acute cannabinoid exposure largely depend on the brain area. For example, a single exposure to THC (3 mg/kg; 15-20 h before) ablated eCB-mediated synaptic plasticity in adult mice NAc and hippocampus (Mato et al., 2004) but not hippocampal CA1 LTP (10 mg/kg; 24 h before) (Hoffman et al., 2007) or eCB-LTD at VTA GABA synapses (Friend et al., 2017). In rats, an acute single injection of WIN (1.2 mg/kg; 24 h before) impaired LTP in the ventral subiculum-accumbens pathway (Abush and Akirav, 2012) and in the Schaffer collateral-CA1 projection (WIN 0.5 mg/kg; 30 min before) (Abush and Akirav, 2010). In a dose-dependent manner, WIN (0.5-2 mg/kg, i.p.) impaired short-term plasticity and long-term potentiation at perforant path dentate gyrus synapses in adult rats (Colangeli et al., 2017). It is important to highlight that in the aforementioned studies only male rodents were evaluated.

Here, we showed that PFC eCB-LTD was ablated in female rats 24 h after SCE regardless of the age, but only adult females had altered neuronal excitability. In contrast, male rats of both ages showed normal eCB-LTD. The eCB signaling machinery is positioned in a way to influence PFC communication and control other brain regions (Hill et al., 2007; Domenici et al., 2006). Sex differences in the eCB system may be involved in these effects. Peak levels of CB1R expression are reached around mid-adolescence in rats (i.e., PND 34-46), and although a higher density of CB1R has been shown in males, a higher G-protein activation after CB1R stimulation is found in adolescent females in several brain areas (Rubino et al., 2008; Burston et al., 2010).

In rodents, the eCB system sexual differences appear early in development (Craft et al., 2013). Sexually dimorphic regulation of synaptic plasticity or intrinsic neuronal activity in the amygdala (Bender et al., 2017; Chen et al., 2014; Fendt et al., 2013), hippocampus (Qi et al., 2016; Harte-Hargrove et al., 2015; Inoue et al., 2014; Huang and Woolley, 2012) and PFC (Li et al., 2016; Nakajima et al., 2014) have been described, but the effects of exogenous cannabinoids on synaptic plasticity in females as well as putative sex differences in its expression remain poorly explored. Our results showed that while eCB-LTD was not affected by cannabinoid exposure in pubescent and adult males, females’ eCB-LTD was ablated. Multiple molecular mechanisms may help explain the observed sex differences. Female rats exhibit greater concentrations of the metabolic enzymes monoacylglycerol lipase (MAGL) and fatty acid amide hydrolase (FAAH) as early as PND 4 compared to males and WIN administration prevents the augmented cell proliferation observed in the amygdala of these animals when compared to males (Krebs-Kraft et al., 2010). Moreover, CB1R expression reaches its peak earlier in females (PND 30) than in males (PND 40) (Romero et al., 1997), whereas at adulthood, CB1R density is lower in the PFC and amygdala of cycling females (Castelli et al., 2014).

While eCB-LTD in both pubescent and adult males was unaffected by SCE, NMDAR-LTP was selectively ablated in adult males. Pubescent males and females of both ages were spared. The eCB system controls NMDAR activity through mecanisms involving signaling pathways and/or direct physical coupling between CB1R and NMDAR NR1 subunits (Rodríguez-Muñoz et al., 2016). Additionally, gonadal hormonal status influences both LTP induction and NMDAR function in male rats (Moradpour et al., 2013). Thus, SCE causes similar behavioral deficits in both male and female rats but triggered different alterations of PFC synapses.

Together, our results reveal behavioral and synaptic sex differences in response to a single in-vivo exposure to cannabinoid. Further analyses of both electrophysiological function and its molecular underpinnings associated with the heightened sensitivity of females to a single in-vivo exposure to cannabinoid may reveal long-term consequences of these early life drug-induced alterations.

## Author contributions

MB, AM, AB, APA and OJM designed research; MB, AM, AB and OL performed research; MB analyzed data; MB, APA and OJM wrote the paper; OJM and APA supervised the entire project. The authors declare no conflict of interest.

## Acknowledgements

The authors are grateful to Dr. Pascale Chavis for helpful discussions, Dr. Andrew Scheyer for critical reading of the manuscript and to the National Institute of Mental Health’s Chemical Synthesis and Drug Supply Program (Rockville, MD, USA) for providing CNQX.

